# TagSeq for gene expression in non-model plants: a pilot study at the Santa Rita Experimental Range NEON core site

**DOI:** 10.1101/2020.04.04.025791

**Authors:** Hannah E. Marx, Stephen Scheidt, Michael S. Barker, Katrina M. Dlugosch

## Abstract

**Premise of the study:** TagSeq is a cost-effective approach for gene expression studies requiring a large number of samples. To date, TagSeq studies in plants have been limited to those with a high quality reference genome. We tested the suitability of reference transcriptomes for TagSeq in non-model plants, as part of a study of natural gene expression variation at the Santa Rita Experimental Range NEON core site.

**Methods:** Tissue for TagSeq was sampled from multiple individuals of four species [*Bouteloua aristidoides* and *Eragrostis lehmanniana* (Poaceae); *Tidestromia lanuginosa* (Amaranthaceae), and *Parkinsonia florida* (Fabaceae)] at two locations on three dates (56 samples total). One sample per species was used to create a reference transcriptome via standard RNA-seq. TagSeq performance was assessed by recovery of reference loci, specificity of tag alignments, and variation among samples.

**Results:** A high fraction of tags aligned to each reference and mapped uniquely. Expression patterns were quantifiable for tens of thousands of loci, which revealed consistent spatial differentiation in expression for all species.

**Discussion:** TagSeq using *de novo* reference transcriptomes was an effective approach to quantifying gene expression in this study. Tags were highly locus specific and generated biologically informative profiles for four non-model plant species.

## INTRODUCTION

Gene expression studies that involve sampling many individuals or tissues can be powerful for identifying variation in transcriptional activity and function (e.g. among populations, over time, in response to the environment or other treatments; Gould et al., 2018; Mead et al., 2019), as well as the structure of transcriptional networks and the genetic basis of gene expression variation (Wisecaver et al., 2017; Li et al., 2020). Such studies are of high interest for non-model species responding to natural environments, as well as for model species (Matz, 2018; Zaidem et al., 2019). Quantitative next generation sequencing of expressed genes, RNA-seq, has made expression studies broadly accessible for non-model species, but remains expensive per sample and difficult to scale up for questions that require high numbers of replicates (Lohman et al., 2016).

A cost-effective approach is to target only a small region of each transcript for sequencing, identifying and quantifying its expression while avoiding sequencing across its full length. Several versions of this approach have involved reading a short tag of sequence upstream of the polyA tail of mRNA. These methods have their roots in one of the first approaches to RNA sequencing prior to the next-generation sequencing era, ‘Expressed Sequence Tags’ or ‘ESTs’ (Parkinson and Blaxter, 2009). Meyer and colleagues (2011) developed an updated version for next generation applications, which continues to be used and adapted (e.g. Dixon et al., 2018; Kremling et al., 2018; Mitchell et al., 2019; Pallares et al., 2020). Recently, Lohman and colleagues (2016) published further developments of what has become known as ‘TagSeq’ (also ‘TAGseq’ or ‘Tag-seq’), and compared its performance with standard RNA-seq of full transcripts. Notably, they find that TagSeq achieves higher accuracy than standard RNA-seq, presumably because sequencing effort is distributed more evenly to all transcripts when only the tag sequence is targeted.

TagSeq tags are short sequences, however, and must be aligned to a reference to fully identify the loci that are being expressed (Meyer et al., 2011). For non-model species and multi-species studies, high quality reference genomes are not likely to be available. Instead, assembled reference transcriptomes can be generated using standard RNA-seq (Matz, 2018). Reference transcriptomes will differ from genomes in that not all loci will be represented by transcripts present in a given sample, not all transcripts will be assembled to full length, and the assembly will vary in the degree to which splice variants, alleles, and paralogs will occur as unique sequences or be merged (Meyer et al., 2011; Yang and Smith, 2013; Carpenter et al., 2019; Patterson et al., 2019). These issues will reduce the number of TagSeq reads that can be uniquely mapped to the reference, relative to a full genome, and they may be particularly problematic in plants where gene and genome duplications are common (Barker et al., 2016; One Thousand Plant Transcriptomes Initiative, 2019; Li and Barker, 2020), though sequencing at the variable 3’ UTR should maximize locus discrimination (Rise et al., 2004).

Meyer at al. (2011) originally demonstrated the TagSeq method in a non-model species of coral, where tags were aligned to a reference transcriptome. Many subsequent studies have successfully used a similar approach in other non-model animals (e.g. Kenkel and Matz, 2016; Dixon et al., 2018; Kriefall et al., 2018; Rocker et al., 2019). In plants, however, TagSeq studies to date appear to have been confined to model species for which a high quality reference genome is available (Meyer et al., 2014; Des Marais et al., 2015; Lovell et al., 2016; Kremling et al., 2018; Chu et al., 2019; Razzaque et al., 2019; Weng et al., 2019). How TagSeq will perform using a reference transcriptome in plants is not clear given the lack of such studies and a paucity of relevant performance information for TagSeq.

Here we report a pilot study using TagSeq to quantify gene expression for four plant species, as part of a study of gene expression variation at the Santa Rita Experimental Range and Core NEON site (Green Valley, AZ, USA). We assembled a reference transcriptome for each species using standard RNA-seq, and analyzed gene expression using TagSeq across multiple individuals for each species, sampled at two locations and three time points. We evaluated the fraction of tags that map uniquely to loci in the reference transcriptome, and the specificity of mapping against references from the same sample, from another sample of the same species, and from other species. We further evaluated the performance of TagSeq in terms of the number of reference loci observed as a function of TagSeq sequencing effort, and the variation in TagSeq profiles across species, sites, and times. Our goal was to assess whether TagSeq is a locus-specific and biologically informative approach for non-model species lacking a high quality reference genome.

## METHODS

### Sampling

Our pilot study focused on four commonly-occurring species at the Santa Rita Experimental Range Long Term Research and Core NEON site (SRER; Appendix 1). These include the native species *Tidestromia lanuginosa* (Nutt.) Standl. (Amaranthaceae; ‘woolly tidestromia’), *Parkinsonia florida* (Benth. ex A. Gray) S. Watson. (Fabaceae; ‘blue palo verde’), and *Bouteloua aristidoides* (Kunth) Griseb. (Poaceae; ‘needle grama’), as well as the introduced species *Eragrostis lehmanniana* Nees (Poaceae; ‘Lehmann lovegrass’; native to southern Africa). All species were identified using a combination of the historical flora of the Santa Rita Experimental Range (Medina, 2003), the Arizona Flora (Kearney et al., 1960), and the Flora of North America (Flora of North America Editorial Committee, eds. 1993). Vouchers were deposited in the University of Arizona herbarium (ARIZ; Appendix 1).

Tissue from mature plants was collected from an apparently healthy individual representing each target species weekly on three dates (Sep 1, 7, and 13) during the 2017 growing season. An entire stem was sampled for *B. aristidoides* (with flowers and fruits) and *E. lehmanniana* (without flowers or fruits). Only leaves and leaflets were sampled for *P. florida* and *T. lanuginosa*. At each sampling date, 2–4 individuals were sampled from each species at each of two locations (4–6 samples total / species / date; Appendix 1). For *P. florida*, samples at the same location and date were not from different individuals, but instead were multiple collections of tissue from the same individual (replicates). The same individual was also resampled at each time point for *P. florida*, and individuals from the same population were sampled for *B. aristidoides, E. lehmanniana*, and *T. lanuginosa*. Samples were collected in the same order on each day beginning at the Phone Pole location and as close as possible to the same time of day (afternoon). Leaf tissues were flash frozen in liquid nitrogen in the field, and transported to the University of Arizona for RNA extraction. Total RNA was extracted from leaf tissue using the Spectrum Plant Total RNA Kit (Sigma-Aldrich, St. Louis, MO, USA) following Protocol A.

The locations included a relatively undisturbed grassland dominated by native species (‘Grassland’), and a more frequently disturbed location near research facilities (‘Phone Pole’). The Grassland location spanned ∼500m along the south side of access Road 424 and was dominated by mostly native grasses (including *Bouteloua aristidoides, Bouteloua barbata var. Rothrockii*, and *Bouteloua repens*) and ocotillo (*Fouquieria splendens*). The Phone Pole location followed a wash along the west side of Road 401 for ∼300m and was dominated by cacti (*Ferocactus wislizeni, Opuntia engelmannii, Cylindropuntia fulgida*), mesquite (*Prosopis velutina*), and creosote (*Larrea tridentata*). The Grassland was roughly 200m higher in elevation than the Phone Pole location and typically receives greater annual moisture (M. McClaran, personal communication).

### RNA-seq for reference transcriptomes

RNA-seq libraries were prepared and sequenced at the Arizona State University’s Biodesign Institute Genomics core facility. Total RNA was used to prepare cDNA using the Ovation RNA-Seq System via single primer isothermal amplification (#7102-A01; Nugen, Redwood CIty, CA, USA) and automated on the Apollo 324 liquid handler (Takara Bio, Kusatsu, Shiga, Japan). cDNA was quantified on the Nanodrop (Thermo Fisher Scientific, Waltham, MA, USA) and was sheared to approximately 300 bp fragments using the Covaris M220 ultrasonicator (Woburn, MA, USA). Libraries were generated using Kapa Biosystem’s library preparation kit (#KK8201; Roche, Basel, Switzerland). Fragments were end repaired and A-tailed, and individual indexes and adapters (#520999; Bioo Scientific, Austin, TX, USA) were ligated onto each sample. The adapter ligated molecules were cleaned using AMPure beads (#A63883; Beckman Coulter, Brea, CA), and amplified with Kapa’s HIFI enzyme (#KK2502). Each library was then analyzed for fragment size on an Agilent Tapestation (Santa Clara, CA, USA), and quantified by qPCR (KAPA Library Quantification Kit #KK4835) on Thermo Fisher Scientific’s Quantstudio 5 before multiplex pooling (13-16 samples per lane in equal representation) and sequencing on the NextSeq500 platform (Illumina, San Diego, California, USA) paired-end 2×150 bp High Output.

Raw reads were filtered and trimmed for adapters and low quality bases using SnoWhite v.2.0.3 (Dlugosch et al., 2013), including TagDust filtering (-D; Lassmann et al., 2009), SeqClean filtering and trimming (-L; Chen et al., 2007)), and a minimum phred score (-Q) of 20. The cleaned read pairs were realigned using fastq-pair (Edwards and Edwards, 2019). Transcripts were assembled using SOAPdenovo-Trans (Xie et al., 2014), using an optimized kmer of 57 (Marx et al., 2020) and archived at https://doi.org/10.5281/zenodo.3740232).

Several aspects of reference assembly quality were assessed. Summary statistics including the number of contig scaffolds, scaffold lengths, and N50 were calculated by Transrate v1.0.3 (Smith-Unna et al., 2016). The completeness of transcriptome coverage was quantified using BUSCO v4.0.5 (Seppey et al., 2019), which identifies representation of a collection of universal single copy orthologs for the viridiplantae (Viridiplantae Odb10) and the eukaryotes (Eukaryote Odb10). Finally, the number of reference contigs matching known proteins were identified using TransPipe (Barker et al., 2010), in which contigs were compared to protein sequences from 25 sequenced and annotated plant genomes from Phytozome (Goodstein et al., 2012) using BLASTX (Wheeler et al., 2008). Best hit proteins were paired with each gene at a minimum cutoff of 30% sequence similarity over at least 150 sites. To determine the reading frame and generate estimated amino acid sequences, each gene was aligned against its best hit protein by Genewise 2.2.2 (Birney et al., 2004). Based on the highest scoring Genewise DNA-protein alignments, stop and ‘N’ containing codons were removed to produce estimated amino acid sequences for each gene (archived at https://doi.org/10.5281/zenodo.3740232).

### TagSeq gene expression

TagSeq libraries were prepared and sequenced at the University of Arizona Genomics core center. Total RNA was used to prepare TagSeq libraries according to the detailed protocol given in Lohman et al. (2016). Specifically, the DNAse I step was included (Qiagen #79254). RNA was fragmented using the NEB Next RNA fragmentation buffer, cleaned using RNA XP Beads (Agencourt), and quantified using RNA PicoGreen (Life Technologies, Carlsbad, California, USA). cDNA was synthesized using forward primers with four degenerate bases near the 3’ end (Eurofins), for the identification of PCR duplicates. cDNA was PCR amplified for 15 cycles, incorporating sample-specific barcodes. PCR products were purified using Ampure XP (Agencourt) beads), and a Pippen Prep was used for 400-500bp size selection. DNA was quantified using DNA PicoGreen (Life Technologies) and pooled in equal representation. The final library was quantified using qPCR (Kapa Biosystems). A total of 56 samples (Appendix 1) were sequenced together on one lane of 1×75bp NextSeq500 High Output (Illumina).

Tag sequences were cleaned of several potential contaminants before analysis. PCR duplicates were identified as sequences that were identical over the first 57 bases, which included the 4 base degenerate primer region, 3 base GGG RNA priming region, and 50 additional bases of unique sequence (using script ‘removePCRdups57.pl’, available at https://doi.org/10.5281/zenodo.3740232). The program ‘cutadapt’ v.1.9.1 (Martin, 2011) was used to trim the 5’ degenerate primer region, 3’ polyA tails (8 or more bases), 3’ low quality bases (min score 20), and primer/adapter contaminants with a minimum overlap of 8bp. Reads less than 57 bases after trimming were discarded. The remaining reads were considered unique sequence tags.

Tags were aligned to the reference transcriptomes using BWA-mem v.0.7.17 (Li and Durbin, 2010) with a bandwidth of 5 bp (-w 5; because gaps relative to the transcriptome reference are not expected in these tag sequences). All other parameters were default. The number of hits to a reference sequence were tallied using HtSeq-count v.0.5.4 (Anders et al., 2015), with --stranded=no (the reference assembly is not stranded). A GTF file was generated from the transcriptome assembly for use with HtSeq-count (using script ‘create_GTF.pl’ available at https://doi.org/10.5281/zenodo.3740232). Hits to each locus were combined across samples and filtered for loci with a minimum of five hits across each species’ dataset, to reduce erroneous hits due to sequencing errors (using script ‘combine_HtSeq.pl’, available athttps://doi.org/10.5281/zenodo.3740232).

We examined the performance of our TagSeq data in terms of recovery of reference loci, specificity of tag alignments, and variation among samples. To assess the ability of TagSeq to track loci in a reference transcriptome, we plotted the proportion of the reference sequences to which tags aligned as a function of TagSeq sequencing effort (total reads), and fitted a logarithmic curve to identify patterns of saturation with sequencing effort. To examine the specificity of tags, we quantified the fraction of tags that mapped to multiple reference loci. We also compared the number of tags aligning to references when i) the reference and TagSeq were derived from the same sample, ii) when the reference and tags were derived from different samples of the same species (individuals or populations), and iii) when the reference and tags were derived from different species.

Finally, we assessed differences among samples with MDS ordination of all TagSeq samples for a species. R/vegan v.2.4-3 (Oksanen et al., 2016) was used to calculate the relative abundance matrix across loci and samples. R/limma v.3.26.9 (Ritchie et al., 2015) was used to calculate root-mean-square deviation (Euclidean distance) among samples and construct the ordination. Distances were based on the loci with the largest standard deviation among all samples (gene.selection = “common”). The number of top loci used was determined by the median value of loci observed among samples: 60k for *B. aristidoides*, 54k for *E. lehmanniana*, 52k for *P. florida*, and 110k for *T. lanuginosa*.

## RESULTS

Raw data for RNA-seq references and TagSeq gene expression were deposited at the NCBI Short Read Archive under BioProject #PRJNA599443. Reference sequencing included 64-83 million raw reads and 63-81 million clean reads, per species (Table 1). Assembly metrics indicated that the most complete assembly was obtained for *P. florida*, with 78% BUSCO recovery, N50 of 895bp, and the largest fraction of contigs translating to known proteins (Table 1). The two grass species yielded the least comprehensive reference assemblies, with 45% and 49% BUSCO scores for *E. lehmanniana* and *B. aristidoides* respectively, and N50 values below 600bp for both species. Assembly metrics were generally intermediate for *T. lanuginosa*, though it had the largest number of assembled contigs and contigs translating to proteins. Notably, despite having the second largest sequencing effort, *E. lehmanniana* had the smallest maximum contig size, lowest BUSCO score, and fewest contigs matching known proteins, suggesting that contigs assembled more poorly for this species relative to the others.

**TABLE 1.**
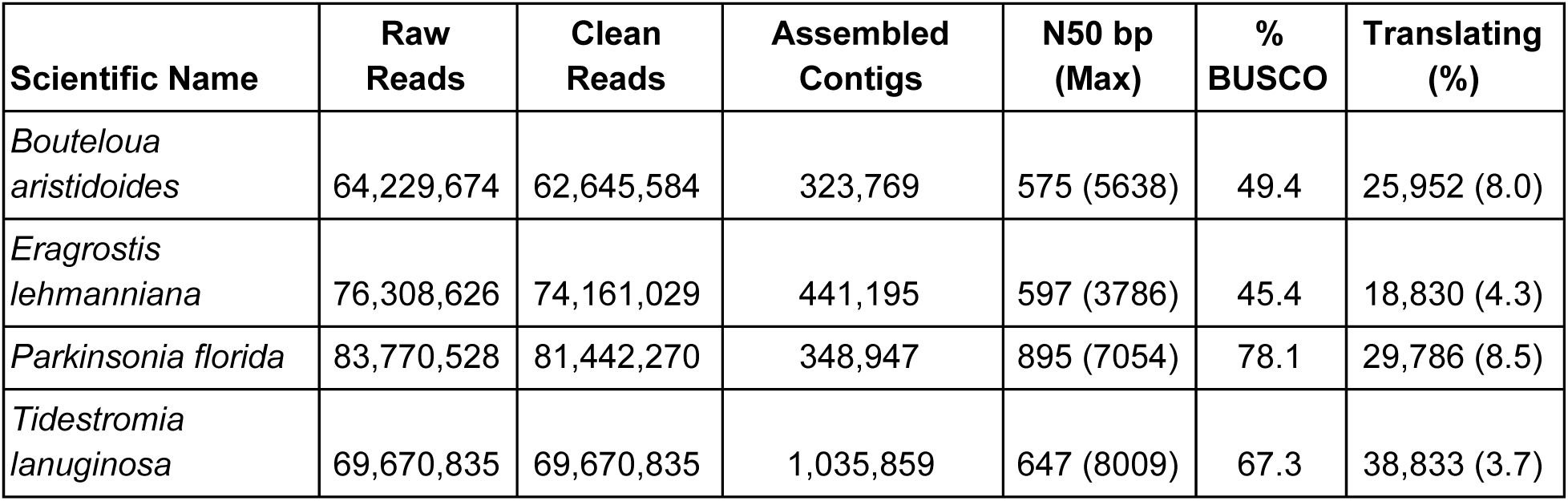
RNAseq reference assembly summary statistics for each species. Included are the numbers of raw reads, clean reads, assembled contigs, and contigs aligning to proteins (translating), as well as the N50 and maximum contig length (bp) and the % of BUSCO sequences matching contigs (complete and partial) in the Viridiplantae database.

| Scientific Name | Raw Reads | Clean Reads | Assembled Contigs | N50 bp (Max) | % BUSCO | Translating (%) |
| --- | --- | --- | --- | --- | --- | --- |
| *Bouteloua aristidoides* | 64,229,674 | 62,645,584 | 323,769 | 575 (5638) | 49.4 | 25,952 (8.0) |
| *Eragrostis lehmanniana* | 76,308,626 | 74,161,029 | 441,195 | 597 (3786) | 45.4 | 18,830 (4.3) |
| *Parkinsonia florida* | 83,770,528 | 81,442,270 | 348,947 | 895 (7054) | 78.1 | 29,786 (8.5) |
| *Tidestromia lanuginosa* | 69,670,835 | 69,670,835 | 1,035,859 | 647 (8009) | 67.3 | 38,833 (3.7) |

TagSeq libraries included a range of 2.6 million to 9.9 million reads per sample, except for two samples with low read count: *E. lehmanniana* Sample 1 with 335k reads, and *T. languinosa* Sample 9 with 1.2 million reads (Appendix 2). Read cleaning resulted in a low proportion of reads removed due to quality issues (typically <10%). In contrast, PCR duplicates accounted for 42-61% of reads (for all samples other than Sample 1, for which PCR duplicates were 71% of reads).

Among the remaining unique tags, >80% of tags aligned to reference sequences for most samples, other than those of *E. lehmanniana*. For *E. lehmanniana*, 56-65% of tags aligned to the reference (Appendix 2). The fraction of tags aligning to more than one reference was low across all samples (<1%). The fraction of the RNA-seq reference sequences that were observed in TagSeq samples ranged from 10-24% (excluding the low read count Samples 1 and 9), resulting in 33-45% of references observed across all samples together. Requiring that a tag be observed at least 5 times reduced the fraction of references observed by approximately half for each species.

The number of reference sequences observed among tag sequences was related to TagSeq sequencing effort (Fig 1, Appendix 2). All species showed trends toward increases in the proportion of reference loci recovered with increasing sequencing effort, though all trends appeared to be saturating and additional sequencing was predicted to result in only modest increases in references observed. TagSeq samples that were the same as the RNA-seq reference sample did not have disproportionately high matches to the reference sequence for their sequencing effort (Fig 1). Tag alignments to the reference sequence were highly species specific, however (Table 2). Between the two grass species (*B. aristidoides* and *E. lehmanniana*), 15-18% of tags aligned to the reference of the other species. For all other combinations of species, 7% or fewer tags aligned to a heterospecific reference.

**TABLE 2.** The proportion of tags from each sample (rows) aligning to each RNAseq reference (columns). Along the diagonal are the proportion aligning to the conspecific reference for the sample, where the reference comes from a different individual (or different tissue collection of the same individual for P. florida) collected at the same location and date. Off the diagonal are alignments of each sample to references from other species.

|  | RNAseq Reference assembly |  |  |  |
| --- | --- | --- | --- | --- |
| TagSeq Sample | *B. aristidoides* | *E. lehmanniana* | *P. florida* | *T. lanuginosa* |
| *B. aristidoides* - 54 | 0.86 | 0.18 | 0.05 | 0.05 |
| *E. lehmanniana* - 49 | 0.15 | 0.65 | 0.03 | 0.03 |
| *P. florida* - 45 | 0.05 | 0.05 | 0.89 | 0.07 |
| *T. lanuginosa* - 25 | 0.03 | 0.03 | 0.03 | 0.85 |

**FIGURE 1.**
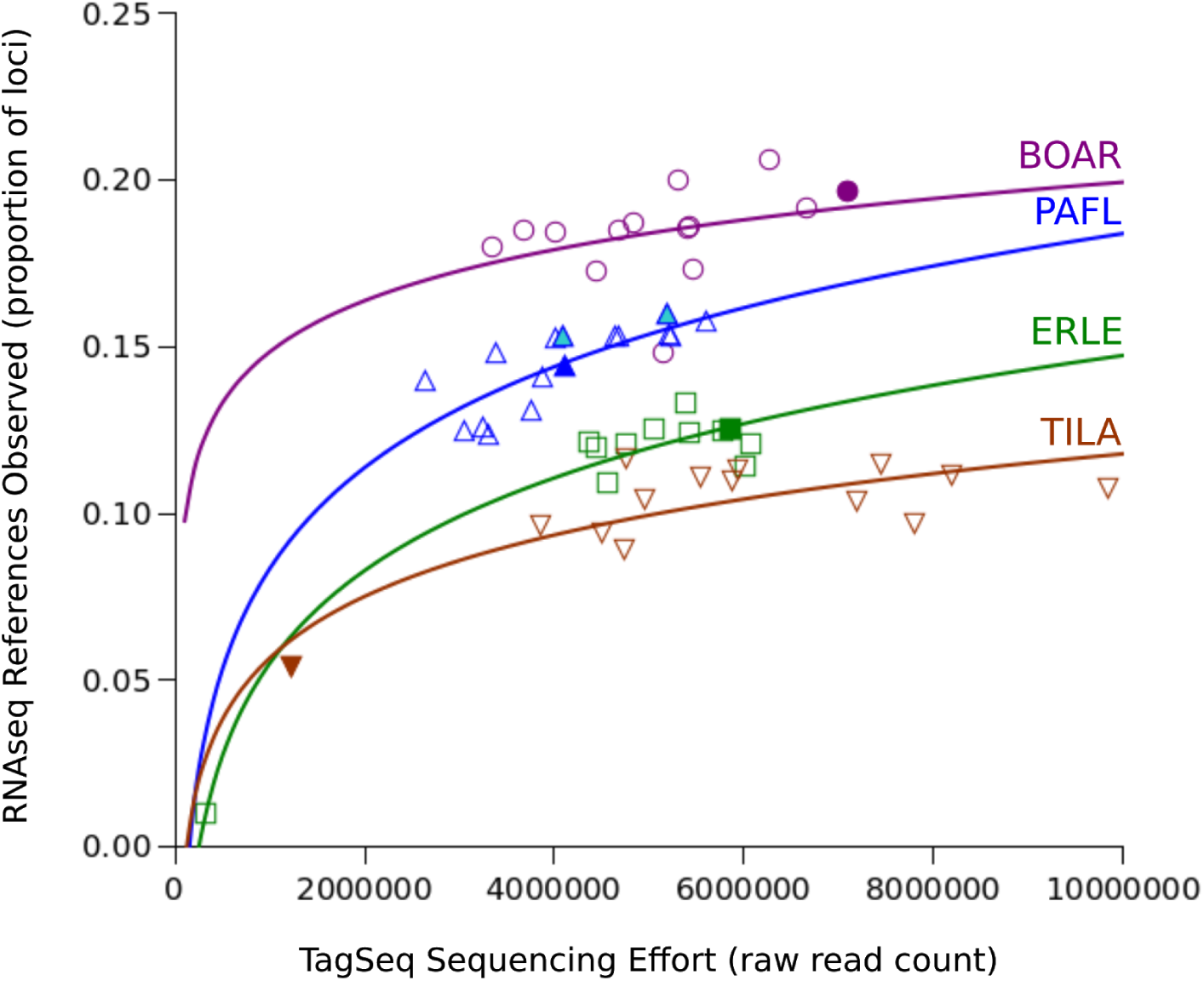
Proportion of RNA-seq reference loci with aligned tags, as a function of sequencing effort (raw read number) of TagSeq libraries. Logarithmic best fits are shown for each species: *B. aristidoides* (BOAR, circles), *E. lehmanniana* (ERLE, squares), *P. florida* (PAFL, upward triangles), and *T. lanuginosa* (TILA, downward triangles). Samples that were used for both a TagSeq library and the RNA-seq reference are indicated by dark filled symbols. Two additional replicates of the reference *P. florida* individual collected on the same day are indicated by lightly shaded symbols. Reference loci were required to be observed in a minimum of 5 tags across a dataset to be counted.

Ordinations for each species revealed clear variation among samples (Fig 2). All species showed clear separation between samples from different locations (closed vs. open symbols, Fig 2). For *E. lehmanniana*, Sample 39 was strongly differentiated from all other samples along Axis 1 (Fig 2B inset), and excluding this sample from the distance matrix allowed further resolution of variation among the remaining samples (Fig 2B). Samples from different dates within a location had a weaker tendency to separate (different symbol shapes; Fig 2), such that samples from the same location and date did not always cluster together.

**FIGURE 2.**
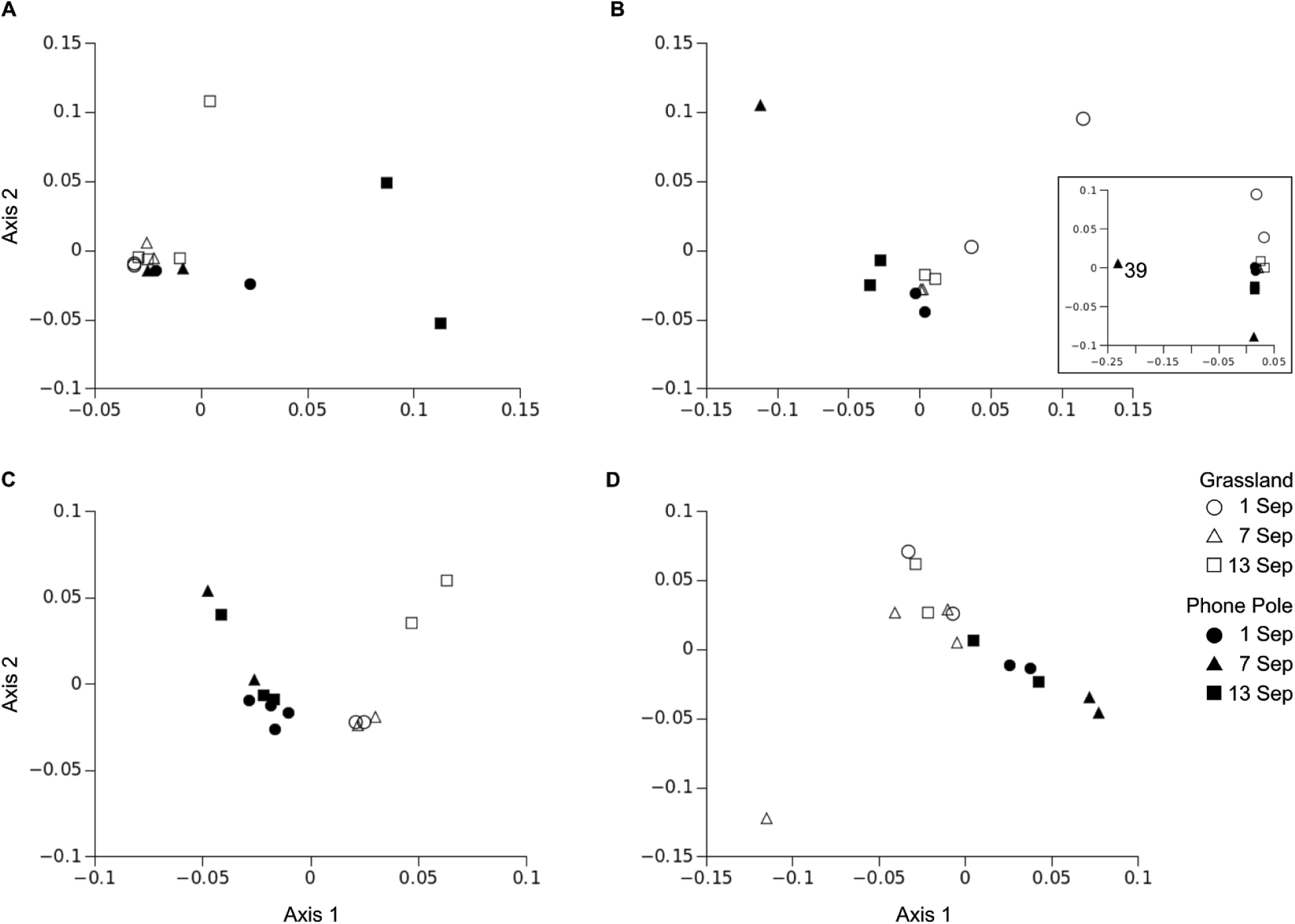
Multidimensional scaling (MDS) ordinations of TagSeq expression data for each species. Tissue samples for *B. aristidoides* (A), *E. lehmanniana* (B), *P. florida* (C), and *T. lanuginosa* (D) were collected from two locations (Grassland, open symbols; Phone Pole, filled symbols) on three dates in 2017 (symbol shapes). Sample 39 for *E. lehmanniana* was highly divergent from others (B, inset) and was removed to better resolve variation among the remaining samples (B).

## DISCUSSION

We evaluated the performance of TagSeq for surveys of gene expression in non-model plant species, using repeated sampling of four species and alignment of tags to *de novo* assemblies of RNA-seq reference transcriptomes. We found that a high fraction of tags aligned to each reference, and few tags mapped to multiple loci or to transcriptomes of other species. Samples from different locations showed clearly differentiated expression profiles for all four species, patterns which were robust to sampling across three dates. Our results support the TagSeq approach as an effective means of generating specific and informative expression profiles in non-model plants.

Quality filtering of tags resulted in very low losses of data (<11% of sequences for all but the most poorly sequenced sample in our dataset), but PCR duplicates comprised 42-60% of samples. PCR duplicates are commonly abundant in Illumina library preparation methods (Aird et al., 2011), and Lohman et al. (2016) reported PCR duplicates of >70% for their test of the protocol used here. The large fraction of sequences involved in PCR duplicates emphasizes the importance of utilizing degenerate bases for identification and removal of duplicates when quantifying expression, as well as the importance of minimizing PCR cycles to maximize sequencing effort on sequences of interest.

For *E. lehmanniana*, >60% of filtered tags mapped to reference loci for most samples, and for nearly all remaining samples of the other species >80% of tags mapped to the reference. For the tags that did not map, at least three factors could explain their failure to align and the variation in alignment rates among species. First, reference loci must include the sequence at the 3’ end of the transcript, immediately upstream of the polyA tail, where TagSeq reads will be located. RNA-seq read distribution is random along the transcript, and therefore many loci will fail to include the required region by chance, and the fraction of loci lacking this region will vary among samples and with the sequencing effort used in creating the reference (Meyer et al., 2011; Conesa et al., 2016; Matz, 2018). Indeed, in particular showed evidence of having the least well-assembled transcriptome among our references. Second, Lohman et al. (2016) found that TagSeq was more sensitive to low levels of expression than was RNA-seq. This difference in sensitivity could result in novel low-expression tags in the TagSeq dataset, for which there is no representative locus in the RNA-seq reference. Finally, allelic differences between samples could cause tags to fail to align to a reference sequence from another individual, though in our dataset we did not see lower rates of alignment in samples that were different than that used for the RNA-seq reference libraries.

For tags lacking a reference sequence, it would be possible in principle to cluster similar tags and to score their expression levels. We observed very low rates (<1%) of mapping to multiple reference loci, which suggests that clustering methods should be able to group tags into inferred loci without high rates of merging across different true loci. Without a reference sequence, however, no information would be available about the identity and function of those loci, which is typically the goal of expression studies (Conesa et al., 2016). Other references (e.g. annotated whole genomes of related species) could be explored for tag identification, but our analyses found that alignment rates to heterospecifics were low (<20% within the same family, <10% between families).

From the perspective of the RNA-seq reference library, a large fraction of reference loci (typically >80%) were not observed in individual TagSeq samples. Again, the samples used for both RNA-seq and TagSeq did not recover a greater number of reference loci, suggesting that neither sequence differences between reference sequences and tags nor differences in genes expressed among samples explained the failure to observe a large number of reference loci in the tags. Additional TagSeq sequencing effort did not result in large gains in the observation of reference loci, though the combination of all samples roughly doubled the fraction of loci observed relative to any one sample, suggesting that tag sequencing effort within the range of our study will affect the number of loci observed. As described above, missing sequence information at the 3’ end of reference loci will also have a large influence on alignment rates, and will set an upper limit on the fraction of loci that can be observed. In their initial publication of the next-generation tag sequencing method, Meyer and colleagues (2011) also report that >80% of reference transcriptome sequences were poorly represented in their tag sequencing, and they suggest that this may be due to sequencing errors in the reference dataset. Only 3.7-8.5% of our reference loci aligned to known proteins, and the number of loci translating to proteins were much more consistent with numbers of genes known from well studied genomes (Marx et al., 2020), suggesting a large number of erroneous loci in our references. These issues regarding reference transcriptome quality could also explain differences in the maximum fraction of loci recovered among the different species.

Finally, we used ordinations to explore whether our resulting TagSeq expression data showed evidence of biologically-relevant structure among samples. Our analyses revealed distinct separation of expression profiles between samples taken from different collection locations within each species. Spatial samples separated into non-overlapping groups along the first (major) axis of ordinations for *P. florida* and *T. lanuginosa*, and through a combination of both axes for *B. aristidoides*. Spatial samples for *E. lehmanniana* converged for a few samples along axis 1. Temporal samples also appeared to group together within spatial locations for some combinations of dates, sites, and species, but additional sampling would be required to resolve temporal patterns robustly. Only one sample (sample 39 for *E. lehmanniana*) across all species was an outlier in ordination space, such that it clustered far from other samples and obscured variation in the remaining data set until it was removed.

In summary, we found that TagSeq expression profiles were biologically informative, and showed little evidence of problems with tag specificity against non-model transcriptome reference datasets. A large proportion of reference loci were not represented in the TagSeq dataset, however, suggesting that completeness of reference assemblies (i.e. assembly of the 3’ end) is likely to influence the identification of loci being expressed. Nevertheless, TagSeq quantified the expression of tens of thousands of loci for each species, and revealed important patterns of differentiation among samples in our dataset.

## ACKNOWLEDGEMENTS

We thank J. Still and J. Galina-Mehlman at the University of Arizona Genomics Core for TagSeq preparation and sequencing, J. Steel and N. Mellor at the Biodesign Institute at Arizona State University for RNA-seq preparation and sequencing, and A. L. Pond, and M. McClaran for field assistance at SRER. This project was supported by NSF #1550838 to M.S.B. and K.M.D., and NSF #1750280 to K.M.D.

## AUTHOR CONTRIBUTIONS

H.E.M., M.S.B, and K.M.D. conceived and designed the experiments. H.E.M. and S.S. collected and geolocated the samples. H.E.M. extracted the RNA and deposited the vouchers. K.M.D. and M.S.B. analyzed the data and drafted the manuscript. All authors contributed to the manuscript revision.

## DATA AVAILABILITY

Raw sequence data for RNA-seq references and TagSeq gene expression have been deposited at the NCBI Short Read Archive under BioProject #PRJNA599443. RNA-seq assemblies, translations, and all custom scripts are available at https://doi.org/10.5281/zenodo.3740232.

## APPENDICES

**APPENDIX 1.** Sample collection site and date, NCBI Short Read Archive (SRA) accession, and information for vouchers deposited at the University of Arizona Herbarium (ARIZ).

| Sample | Site | Latitude | Longitude | Date Collected | SRA Sample Name | ARIZ Catalog# |
| --- | --- | --- | --- | --- | --- | --- |
| *Bouteloua aristidoides* |  |  |  |  |  |  |
| 2* | Native Grassland | 31.8695 | -110.81533 | 13-Sep-17 | Marx 2017-074-a-09132017 |  |
| 6 | Phone Pole | 31.88004 | -110.89882 | 13-Sep-17 | Marx 2017-059-a-09132017 |  |
| 20 | Phone Pole | 31.88004 | -110.89882 | 1-Sep-17 | Marx 2017-003-a-09012017 | 435193 |
| 21 | Phone Pole | 31.88004 | -110.89882 | 7-Sep-17 | Marx 2017-031-a-09072017 |  |
| 23 | Native Grassland | 31.8695 | -110.81533 | 1-Sep-17 | Marx 2017-018-a-09012017 | 435325 |
| 24 | Native Grassland | 31.8695 | -110.81533 | 7-Sep-17 | Marx 2017-046-a-09072017 |  |
| 27 | Native Grassland | 31.8695 | -110.81533 | 1-Sep-17 | Marx 2017-018-b-09012017 | 435325 |
| 28 | Native Grassland | 31.8695 | -110.81533 | 7-Sep-17 | Marx 2017-046-b-09072017 |  |
| 32 | Phone Pole | 31.88004 | -110.89882 | 1-Sep-17 | Marx 2017-003-c-09012017 | 435193 |
| 36 | Phone Pole | 31.88004 | -110.89882 | 13-Sep-17 | Marx 2017-059-c-09132017 |  |
| 41 | Phone Pole | 31.88004 | -110.89882 | 7-Sep-17 | Marx 2017-031-b-09072017 |  |
| 51 | Native Grassland | 31.8695 | -110.81533 | 13-Sep-17 | Marx 2017-074-b-09132017 |  |
| 54 | Native Grassland | 31.8695 | -110.81533 | 13-Sep-17 | Marx 2017-074-c-09132017 |  |
| 56 | Native Grassland | 31.8695 | -110.81533 | 13-Sep-17 | Marx 2017-074-d-09132017 |  |
| *Eragrostis lehmanniana* |  |  |  |  |  |  |
| 1 | Native Grassland | 31.868985 | -110.81573 | 13-Sep-17 | Marx 2017-084-a-09132017 |  |
| 3 | Phone Pole | 31.88004 | -110.89882 | 7-Sep-17 | Marx 2017-032-a-09072017 |  |
| 10* | Native Grassland | 31.868985 | -110.81573 | 1-Sep-17 | Marx 2017-028-a-09012017 | 435319 |
| 15 | Native Grassland | 31.868985 | -110.81573 | 7-Sep-17 | Marx 2017-056-a-09072017 |  |
| 16 | Phone Pole | 31.88004 | -110.89882 | 13-Sep-17 | Marx 2017-060-a-09132017 |  |
| 17 | Phone Pole | 31.88004 | -110.89882 | 1-Sep-17 | Marx 2017-004-a-09012017 | 435194 |
| 26 | Native Grassland | 31.868985 | -110.81573 | 7-Sep-17 | Marx 2017-056-b-09072017 |  |
| 30 | Native Grassland | 31.868985 | -110.81573 | 13-Sep-17 | Marx 2017-084-b-09132017 |  |
| 39 | Phone Pole | 31.88004 | -110.89882 | 7-Sep-17 | Marx 2017-032-b-09072017 |  |
| 43 | Phone Pole | 31.88004 | -110.89882 | 13-Sep-17 | Marx 2017-060-b-09132017 |  |
| 44 | Phone Pole | 31.88004 | -110.89882 | 1-Sep-17 | Marx 2017-004-b-09012017 | 435194 |
| 49 | Native Grassland | 31.868985 | -110.81573 | 1-Sep-17 | Marx 2017-028-b-09012017 | 435319 |
| *Parkinsonia florida* |  |  |  |  |  |  |
| 11 | Phone Pole | 31.880457 | -110.89868 | 1-Sep-17 | Marx 2017-014-a-09012017 | 435322 |
| 12* | Phone Pole | 31.880457 | -110.89868 | 13-Sep-17 | Marx 2017-070-a-09132017 |  |
| 13 | Native Grassland | 31.869053 | -110.81528 | 7-Sep-17 | Marx 2017-051-b-09072017 |  |
| 14 | Native Grassland | 31.869053 | -110.81528 | 1-Sep-17 | Marx 2017-023-b-09012017 | 435326 |
| 19 | Phone Pole | 31.880457 | -110.89868 | 1-Sep-17 | Marx 2017-014-b-09012017 | 435322 |
| 22 | Phone Pole | 31.880457 | -110.89868 | 7-Sep-17 | Marx 2017-042-a-09072017 |  |
| 31 | Native Grassland | 31.869053 | -110.81528 | 7-Sep-17 | Marx 2017-051-c-09072017 |  |
| 34 | Native Grassland | 31.869053 | -110.81528 | 13-Sep-17 | Marx 2017-079-b-09132017 |  |
| 37 | Native Grassland | 31.869053 | -110.81528 | 13-Sep-17 | Marx 2017-079-c-09132017 |  |
| 38 | Phone Pole | 31.880457 | -110.89868 | 13-Sep-17 | Marx 2017-070-b-09132017 |  |
| 40 | Phone Pole | 31.880457 | -110.89868 | 7-Sep-17 | Marx 2017-042-b-09072017 |  |
| 45 | Phone Pole | 31.880457 | -110.89868 | 13-Sep-17 | Marx 2017-070-c-09132017 |  |
| 46 | Phone Pole | 31.880457 | -110.89868 | 1-Sep-17 | Marx 2017-014-c-09012017 | 435322 |
| 50 | Phone Pole | 31.880457 | -110.89868 | 1-Sep-17 | Marx 2017-014-d-09012017 | 435322 |
| 52 | Native Grassland | 31.869053 | -110.81528 | 7-Sep-17 | Marx 2017-051-d-09072017 |  |
| 55 | Native Grassland | 31.869053 | -110.81528 | 1-Sep-17 | Marx 2017-023-d-09012017 | 435326 |
| *Tidestromia lanuginosa* |  |  |  |  |  |  |
| 4 | Phone Pole | 31.879973 | -110.89881 | 1-Sep-17 | Marx 2017-008-a-09012017 | 435324 |
| 5 | Phone Pole | 31.879973 | -110.89881 | 13-Sep-17 | Marx 2017-064-a-09132017 |  |
| 7 | Native Grassland | 31.869432 | -110.81542 | 13-Sep-17 | Marx 2017-072-a-09132017 |  |
| 8 | Phone Pole | 31.879973 | -110.89881 | 7-Sep-17 | Marx 2017-036-a-09072017 |  |
| 9* | Native Grassland | 31.869432 | -110.81542 | 7-Sep-17 | Marx 2017-043-a-09072017 |  |
| 18 | Native Grassland | 31.869432 | -110.81542 | 1-Sep-17 | Marx 2017-016-a-09012017 | 435323 |
| 25 | Native Grassland | 31.869432 | -110.81542 | 7-Sep-17 | Marx 2017-043-b-09072017 |  |
| 29 | Native Grassland | 31.869432 | -110.81542 | 13-Sep-17 | Marx 2017-072-b-09132017 |  |
| 33 | Phone Pole | 31.879973 | -110.89881 | 1-Sep-17 | Marx 2017-008-b-09012017 | 435324 |
| 35 | Phone Pole | 31.879973 | -110.89881 | 7-Sep-17 | Marx 2017-036-b-09072017 |  |
| 42 | Native Grassland | 31.869432 | -110.81542 | 1-Sep-17 | Marx 2017-016-b-09012017 | 435323 |
| 47 | Phone Pole | 31.879973 | -110.89881 | 13-Sep-17 | Marx 2017-064-b-09132017 |  |
| 48 | Native Grassland | 31.869432 | -110.81542 | 7-Sep-17 | Marx 2017-043-c-09072017 |  |
| 53 | Native Grassland | 31.869432 | -110.81542 | 7-Sep-17 | Marx 2017-043-d-09072017 |  |

\*Samples which were used as both RNAseq reference transcriptome libraries and TagSeq expression libraries.

**APPENDIX 2.** TagSeq summary statistics for each sample. Mult. Hits refers to hits to multiple reference loci. Cont./Qual. refers to tags removed due to contamination or low quality scores.

|  | Count: | Count: | Prop. of reads removed due to: |  |  | Count: | Prop. of tags: |  |  | Prop. of ref. contigs: |  |
| --- | --- | --- | --- | --- | --- | --- | --- | --- | --- | --- | --- |
| Sample | Raw Reads | Clean Tags | PCR Duplicates | Short | Cont./Qual. | Tags Aligned | Aligned | Not Aligned | Mult. Hits | Total Hits | Min. 5 Hits |
| *Bouteloua aristidoides* |  |  |  |  |  |  |  |  |  |  |  |
| 2* | 7108820 | 2864252 | 0.54 | 0.0001 | 0.05 | 2420723 | 0.85 | 0.15 | 0.004 | 0.22 | 0.20 |
| 6 | 6659522 | 2617957 | 0.56 | 0.0001 | 0.05 | 2079881 | 0.79 | 0.21 | 0.005 | 0.22 | 0.19 |
| 20 | 5317488 | 2455245 | 0.47 | 0.0001 | 0.06 | 2045210 | 0.83 | 0.17 | 0.004 | 0.23 | 0.20 |
| 21 | 5163416 | 2201758 | 0.52 | 0.0000 | 0.05 | 1096620 | 0.50 | 0.50 | 0.003 | 0.16 | 0.15 |
| 23 | 5464515 | 2214799 | 0.53 | 0.0001 | 0.06 | 1879440 | 0.85 | 0.15 | 0.005 | 0.19 | 0.17 |
| 24 | 4684318 | 2176801 | 0.48 | 0.0001 | 0.06 | 1754713 | 0.81 | 0.19 | 0.004 | 0.20 | 0.18 |
| 27 | 4451466 | 1939050 | 0.50 | 0.0001 | 0.06 | 1643622 | 0.85 | 0.15 | 0.004 | 0.19 | 0.17 |
| 28 | 4849566 | 2143817 | 0.50 | 0.0001 | 0.06 | 1810488 | 0.84 | 0.16 | 0.005 | 0.21 | 0.19 |
| 32 | 5438574 | 2415989 | 0.50 | 0.0001 | 0.05 | 2024038 | 0.84 | 0.16 | 0.004 | 0.20 | 0.19 |
| 36 | 6268774 | 2748050 | 0.50 | 0.0001 | 0.06 | 2284930 | 0.83 | 0.17 | 0.005 | 0.24 | 0.21 |
| 41 | 5405364 | 2246322 | 0.53 | 0.0001 | 0.05 | 1859298 | 0.83 | 0.17 | 0.005 | 0.21 | 0.19 |
| 51 | 3695704 | 1751372 | 0.45 | 0.0001 | 0.07 | 1471954 | 0.84 | 0.16 | 0.005 | 0.21 | 0.18 |
| 54 | 3355992 | 1696597 | 0.42 | 0.0002 | 0.07 | 1455374 | 0.86 | 0.14 | 0.004 | 0.20 | 0.18 |
| 56 | 4009876 | 1900705 | 0.46 | 0.0001 | 0.07 | 1629273 | 0.86 | 0.14 | 0.004 | 0.20 | 0.18 |
| TOTAL |  | 31372714 |  |  |  |  |  |  |  | 0.45 | 0.27 |
| *Eragrostis lehmanniana* |  |  |  |  |  |  |  |  |  |  |  |
| 1 | 335464 | 17904 | 0.71 | 0.0073 | 0.22 | 10362 | 0.58 | 0.42 | 0.006 | 0.01 | 0.01 |
| 3 | 6071239 | 2538484 | 0.53 | 0.0001 | 0.05 | 1592331 | 0.63 | 0.37 | 0.005 | 0.14 | 0.12 |
| 10* | 5866891 | 2366892 | 0.55 | 0.0000 | 0.04 | 1522526 | 0.64 | 0.36 | 0.006 | 0.15 | 0.12 |
| 15 | 6013007 | 2267378 | 0.58 | 0.0000 | 0.04 | 1457729 | 0.64 | 0.36 | 0.005 | 0.13 | 0.11 |
| 16 | 5779005 | 2374148 | 0.54 | 0.0001 | 0.05 | 1542273 | 0.65 | 0.35 | 0.005 | 0.15 | 0.12 |
| 17 | 5440165 | 2303471 | 0.52 | 0.0000 | 0.05 | 1427988 | 0.62 | 0.38 | 0.005 | 0.14 | 0.12 |
| 26 | 4772169 | 2096114 | 0.50 | 0.0001 | 0.06 | 1374445 | 0.66 | 0.34 | 0.005 | 0.14 | 0.12 |
| 30 | 4458162 | 1968870 | 0.50 | 0.0001 | 0.06 | 1293357 | 0.66 | 0.34 | 0.006 | 0.14 | 0.12 |
| 39 | 5386577 | 2705308 | 0.44 | 0.0001 | 0.05 | 1520821 | 0.56 | 0.44 | 0.004 | 0.17 | 0.13 |
| 43 | 5067830 | 2199399 | 0.51 | 0.0001 | 0.06 | 1401928 | 0.64 | 0.36 | 0.005 | 0.15 | 0.13 |
| 44 | 4373470 | 1993560 | 0.49 | 0.0000 | 0.06 | 1209574 | 0.61 | 0.39 | 0.005 | 0.14 | 0.12 |
| 49 | 4559967 | 1803395 | 0.56 | 0.0001 | 0.04 | 1169445 | 0.65 | 0.35 | 0.006 | 0.12 | 0.11 |
| TOTAL |  | 24634923 |  |  |  |  |  |  |  | 0.33 | 0.17 |
| *Parkinsonia florida* |  |  |  |  |  |  |  |  |  |  |  |
| 11 | 3887959 | 1432767 | 0.53 | 0.0003 | 0.10 | 1253182 | 0.87 | 0.13 | 0.005 | 0.15 | 0.14 |
| 12* | 4110903 | 1603854 | 0.55 | 0.0001 | 0.06 | 1444493 | 0.90 | 0.10 | 0.005 | 0.16 | 0.14 |
| 13 | 3305618 | 1146180 | 0.58 | 0.0002 | 0.08 | 992508 | 0.87 | 0.13 | 0.005 | 0.13 | 0.12 |
| 14 | 3066122 | 1155170 | 0.58 | 0.0001 | 0.04 | 982613 | 0.85 | 0.15 | 0.005 | 0.13 | 0.12 |
| 19 | 5207037 | 1779503 | 0.60 | 0.0001 | 0.06 | 1607565 | 0.90 | 0.10 | 0.005 | 0.17 | 0.15 |
| 22 | 4643217 | 1704905 | 0.56 | 0.0001 | 0.07 | 1524181 | 0.89 | 0.11 | 0.005 | 0.17 | 0.15 |
| 31 | 3250480 | 1140899 | 0.59 | 0.0001 | 0.06 | 1010075 | 0.89 | 0.11 | 0.005 | 0.13 | 0.13 |
| 34 | 4694661 | 1931488 | 0.52 | 0.0003 | 0.07 | 1697593 | 0.88 | 0.12 | 0.005 | 0.17 | 0.15 |
| 37 | 5231677 | 2026252 | 0.54 | 0.0002 | 0.07 | 1772085 | 0.87 | 0.13 | 0.005 | 0.17 | 0.15 |
| 38 | 5194831 | 2064010 | 0.53 | 0.0002 | 0.07 | 1864999 | 0.90 | 0.10 | 0.005 | 0.18 | 0.16 |
| 40 | 5607535 | 2098556 | 0.55 | 0.0002 | 0.07 | 1856128 | 0.88 | 0.12 | 0.004 | 0.17 | 0.16 |
| 45 | 4106462 | 1598327 | 0.54 | 0.0001 | 0.07 | 1427567 | 0.89 | 0.11 | 0.005 | 0.17 | 0.15 |
| 46 | 3395667 | 1362410 | 0.52 | 0.0001 | 0.07 | 1224492 | 0.90 | 0.10 | 0.005 | 0.16 | 0.15 |
| 50 | 4011175 | 1550240 | 0.55 | 0.0001 | 0.07 | 1400355 | 0.90 | 0.10 | 0.005 | 0.17 | 0.15 |
| 52 | 2638261 | 1165622 | 0.46 | 0.0005 | 0.09 | 1010315 | 0.87 | 0.13 | 0.005 | 0.16 | 0.14 |
| 55 | 3761388 | 1348379 | 0.56 | 0.0001 | 0.08 | 1168972 | 0.87 | 0.13 | 0.005 | 0.14 | 0.13 |
| TOTAL |  | 25108562 |  |  |  |  |  |  |  | 0.36 | 0.21 |
| *Tidestromia lanuginosa* |  |  |  |  |  |  |  |  |  |  |  |
| 4 | 8206366 | 3314540 | 0.55 | 0.0001 | 0.04 | 2698441 | 0.81 | 0.19 | 0.005 | 0.13 | 0.11 |
| 5 | 7206798 | 2879104 | 0.56 | 0.0000 | 0.04 | 2359448 | 0.82 | 0.18 | 0.005 | 0.12 | 0.10 |
| 7 | 7803507 | 2732931 | 0.61 | 0.0001 | 0.04 | 2266700 | 0.83 | 0.17 | 0.004 | 0.11 | 0.10 |
| 8 | 9851038 | 3308006 | 0.63 | 0.0000 | 0.03 | 2723453 | 0.82 | 0.18 | 0.005 | 0.13 | 0.11 |
| 9* | 1241373 | 555457 | 0.45 | 0.0021 | 0.11 | 436457 | 0.79 | 0.21 | 0.004 | 0.06 | 0.05 |
| 18 | 4744115 | 1982995 | 0.53 | 0.0000 | 0.06 | 1538724 | 0.78 | 0.22 | 0.004 | 0.10 | 0.09 |
| 25 | 5542647 | 2477967 | 0.49 | 0.0001 | 0.06 | 2100990 | 0.85 | 0.15 | 0.005 | 0.14 | 0.11 |
| 29 | 5944645 | 2594919 | 0.52 | 0.0000 | 0.05 | 2155507 | 0.83 | 0.17 | 0.005 | 0.14 | 0.11 |
| 33 | 5891911 | 2708682 | 0.49 | 0.0001 | 0.05 | 2281047 | 0.84 | 0.16 | 0.005 | 0.13 | 0.11 |
| 35 | 7448586 | 3111221 | 0.53 | 0.0001 | 0.05 | 2559379 | 0.82 | 0.18 | 0.004 | 0.14 | 0.11 |
| 42 | 4957036 | 2249422 | 0.49 | 0.0001 | 0.06 | 1812268 | 0.81 | 0.19 | 0.004 | 0.13 | 0.10 |
| 47 | 4508978 | 1871639 | 0.53 | 0.0000 | 0.06 | 1563872 | 0.84 | 0.16 | 0.005 | 0.11 | 0.09 |
| 48 | 3857432 | 1822810 | 0.46 | 0.0001 | 0.07 | 1544443 | 0.85 | 0.15 | 0.004 | 0.11 | 0.10 |
| 53 | 4770648 | 2282278 | 0.47 | 0.0001 | 0.06 | 1884754 | 0.83 | 0.17 | 0.004 | 0.14 | 0.12 |
| TOTAL |  | 33891971 |  |  |  |  |  |  |  | 0.37 | 0.19 |

\*Samples which were used as both RNAseq reference transcriptome libraries and TagSeq expression libraries.

